# Inflated citations and metrics of journals discontinued from Scopus for publication concerns: the GhoS(t)copus Project

**DOI:** 10.1101/2020.03.26.007435

**Authors:** Andrea Cortegiani, Mariachiara Ippolito, Giulia Ingoglia, Andrea Manca, Lucia Cugusi, Anna Severin, Michaela Strinzel, Vera Panzarella, Giuseppina Campisi, Lalu Manoj, Cesare Gregoretti, Sharon Einav, David Moher, Antonino Giarratano

**Author notes:** **Corresponding author**: Andrea Cortegiani. Department of Surgical, Oncological and Oral Science (Di.Chir.On.S). Section of Anaesthesia, Analgesia, Intensive Care and Emergency. Policlinico Paolo Giaccone, University of Palermo, via del Vespro 129, 90127, Palermo, Italy.

## Abstract

**Background:** Scopus is a leading bibliometric database. It contains the largest number of articles cited in peer-reviewed publications. The journals included in Scopus are periodically re-evaluated to ensure they meet indexing criteria and some journals might be discontinued for publication concerns. These journals remain indexed and can be cited. Their metrics have yet to be studied. This study aimed to evaluate the main features and metrics of journals discontinued from Scopus for publication concerns, before and after their discontinuation, and to determine the extent of predatory journals among the discontinued journals.

**Methods:** We surveyed the list of discontinued journals from Scopus (July 2019). Data regarding metrics, citations and indexing were extracted from Scopus or other scientific databases, for the journals discontinued for publication concerns.

**Results:** A total of 317 journals were evaluated. Ninety-three percent of the journals (294/318) declared they published using an Open Access model. The subject areas with the greatest number of discontinued journals were *Medicine* (52/317; 16%), *Agriculture and Biological Science* (34/317; 11%), and *Pharmacology, Toxicology and Pharmaceutics* (31/317; 10%). The mean number of citations per year after discontinuation was significantly higher than before (median of difference 64 citations, p<0.0001), and so was the number of citations per document (median of difference 0.4 citations, p<0.0001). Twenty-two percent (72/317) were included in the Cabell’s blacklist. The DOAJ currently included only 9 journals while 61 were previously included and discontinued, most for “suspected editorial misconduct by the publisher’. **Conclusions:** The citation count of journals discontinued for publication concerns increases despite discontinuation and predatory behaviors seemed common. This paradoxical trend can inflate scholars’ metrics prompting artificial career advancements, bonus systems and promotion. Countermeasures should be taken urgently to ensure the reliability of Scopus metrics both at the journal- and author-level for the purpose of scientific assessment of scholarly publishing.

## Introduction

Scopus is a leading bibliometric database launched in 2004 by the publishing and analytics company Elsevier. It was developed by research institutions, researchers and librarians, and contains the largest number of abstracts and articles cited in peer reviewed academic journal articles that cover scientific, technical, medical, and social science fields.[1]

Scopus provides bibliometric indicators that many institutions use to rank journals to evaluate the track record of scholars who seek hiring or promotion. These metrics are also used to allocate financial bonuses or to evaluate funding applications. [2,3,4] Ensuring the quality of the content of the Scopus database is, therefore, of great importance.

To be indexed in Scopus, journals are evaluated and periodically reviewed by an independent and international Content Selection and Advisory Board (CSAB), which is a group of scientists, researchers and librarians, comprised of 17 Subject Chairs, each representing a specific subject field, and by a computerized algorithm. [1] At any time after a journal inclusion, concerns regarding its quality may be raised by a formal complaint, thereby flagging the journal for re-evaluation by the CSAB. Should the CSAB panel determine that the journal no longer meets Scopus standards, new articles from that journal are no longer be indexed. [1] One of the most common reasons for discontinuation is ‘publication concerns’, which refers to the quality of editorial practices or other issues that have an impact on its suitability for continued coverage. [5] The list of the discontinued sources is publicly available and is updated approximately every six months. [6] However, publications from no longer indexed journals may not be removed retrospectively from Scopus. Hence, articles indexed prior to the date of discontinuation could remain part of the database. [7]

It has been claimed that a number of journals discontinued from Scopus for publication concerns might be so-called ‘predatory’ journals. [6,7,8] Predatory journals “prioritize self-interest at the expense of scholarship and are characterized by false or misleading information, deviation from best editorial and publication practices, a lack of transparency, and/or the use of aggressive and indiscriminate solicitation practices”. [9] Since researchers are pressured to publish in indexed journals, predatory journals are constantly trying to be indexed in the Scopus database, thereby boosting their attractiveness to researchers. [2,7,8,10] Having articles from predatory journals indexed in Scopus poses a threat to the credibility of science and might cause harm particularly in fields where practitioners rely on empirical evidence in the form of indexed journal articles. [10,11]

We hypothesize that, even though Scopus coverage is halted for discontinued journals, still they can get citations, as all their documents already indexed remain available to users. To date, the metrics of those journals discontinued for publication concerns have not been studied yet. Therefore, by the present analysis we set out to (1) evaluate the main scientific features and citation metrics of journals discontinued from Scopus for publication concerns, before and after discontinuation, and (2) determine the extent of predatory journals included in the discontinued journals.

## Methods

### Search strategy

The freely accessible and regularly updated Elsevier list [1] of journals discontinued from the Scopus database (version July 2019) [12] was accessed on 24^th^ January 2020 (See **S1 Appendix)**. We restricted our analysis to journals discontinued for “publication concerns”. Journals were checked for relevant data (described below), then independently collected by eight of the authors in pairs (MI, GI, AM, LC, AS, MS, VP, AC) using a standardized data extraction form. A second check was performed by other four authors (LM, CG, SE, AG) to confirm the data and resolve discrepancies. Data collection was initiated on 24^th^ January and completed by the end of February 2020. Confirmed data were registered on an Excel datasheet (**S2 Appendix**).

### Retrieved data and sources

Data were extracted either from the Scopus database [12] or by searching other sources, such as SCImago Journal & Country Rank (SJCR), [13] Journal Citation Reports, Centre for Science and Technology Studies (CWTS) Journal Indicators, [14] Beall’s updated List, [15] Directory of Open Access Journals (DOAJ), [16] PubMed [17] and Web of Science. [18] Open Access policy was checked directly on journals websites. A standardized data extraction form, independently applied by eight authors (MI, GI, AM, LC, AS, MS, VP, AC), was used to collect the following data: journal title, name and country of the publisher, the number of years of Scopus coverage, year of Scopus discontinuation, subject areas and sub-subject areas, Impact Factor (IF), CiteScore, SCImago Journal Rank (SJR), Source Normalized Impact *per* Paper (SNIP), best SCImago quartile, inclusion in PubMed, Web Of Science (WOS) and DOAJ (for open access journals) databases, presence in the updated Beall’s List, total number of published documents and total number of citations. All the metrics were checked on the year of Scopus discontinuation. In cases of discrepancies between Scopus data and other sources, the Scopus database was used as the preferential source.

We defined the ‘before discontinuation’ time frame as the period comprised within the first year of journal coverage by Scopus and the year of discontinuation, which was not included in our calculations. By ‘after discontinuation’ time frame, we referred to the period comprised within the year of Scopus discontinuation and the year of our data collection. In cases of multiple discontinuations, we considered only the last one, according to the date of the last document displayed in the Scopus database. Citations ‘before’ and ‘after’ the date of discontinuation were manually counted based on either the Scopus journal overview or the downloadable tables made available by Scopus upon request. When evaluating the journal inclusion in PubMed, WOS and DOAJ, year 2019 was considered as the reference year, thus preventing any disadvantage for journals with a time gap for publication.

Finally, one author (AS) checked whether discontinued journals were present in Cabell’s whitelist or blacklist [19] or the DOAJ’s list of discontinued journals. [20] As some of the journals included in the blacklist lack ISSNs or other unique identifiers, the comparison of the three lists with Scopus’s discontinued journals was based on matching the journals’ names by similarity using the Jaro-Winkler algorithm in R package RecordLinkage, following the approach developed by Strinzel et al. (2019). [21,22] The Jaro-Winkler metric, scaled between 0 (no similarity) and 1 (exact match), was calculated for all possible journals’ pairings. [23] We manually inspected all pairs with a Jaro-Winkler metric smaller than one in order to include cases where, due to the orthographical differences between the lists, no exact match was found. For each matched pair, we compared the journals’ publishers and, where possible, ISSNs, to exclude any cases where two journals had the same or a similar name but were edited by different publishers.

Full definitions and descriptions of the sources and metrics are reported in the **S3 Appendix**.

### Statistical analysis

All data management and calculations were performed using Microsoft Excel (version 2013, Microsoft Corporation®, USA) and GraphPad Prism (version 8.3.1, 322, GraphPad software®, San Diego California). The normality of the distribution was assessed with the D’Agostino-Pearson test. Means and standard deviations (SDs) for variables with a normal distribution or medians, interquartile ranges (IQRs, 25th– 75th) and ranges (minimum value - maximum value) for non-normally distributed data were calculated and reported. Categorical data were expressed as proportions and percentages.

The paired sample *t* test or the Wilcoxon matched-pairs signed ranked test were used to compare journals’ data before and after Scopus discontinuation, as appropriate.

## Results

Data could be retrieved regarding 317 of the 348 journals listed as discontinued (91.1%).

### Journals’ and publishers’ characteristics

Among the 135 publishers identified, the publishers with the largest number of discontinued journals were: *Academic Journals Inc*. (39/317; 12.3%), *Asian Network for Scientific Information* (19/317; 6.0%), and *OMICS Publishing Group* (18/317; 5.7%). **S1 Table** reports the distribution of Scopus discontinued journals by publisher. United States (76/317, 24%), India (63/317, 20%) and Pakistan (49/317, 15%) were the most common countries where publishers declared they were headquartered (**S1 Fig**. and **S2 Table**).

The subject areas with the greatest number of discontinued journals were *Medicine* (52/317; 16%), *Agriculture and Biological Science* (34/317; 11%), and *Pharmacology, Toxicology and Pharmaceutics* (31/317; 10%) **S3 Table** and **S4 Table** report the distribution of discontinued journals by subject area and sub-area in full. Ninety-three percent of the journals (294/318) declared they published using an Open Access model.

**Table 1** shows the characteristics and metrics of journals at the time of discontinuation. The median time of Scopus coverage prior to discontinuation of the journals was 8 years (IQR 6-10, range 1-54). Two hundred ninety-nine journals had been assigned to a SCImago quartile (Q); 39 of them (13%) listed in Q1 or Q2, and 260 in Q3 or Q4 (87%). Only ten of the discontinued journals had an Impact Factor at the year of discontinuation, with a median value of 0.84 (IQR 0.37-2.29, range 0.28-4).

**Table 1.** Journals characteristics at the year of Scopus discontinuation.

|  |  |
| --- | --- |
| **Scopus coverage (yrs.) \*** | 8 [6-10] (1-54) |
| **Time from Scopus discontinuation (yrs.) \*** | 5 [4-6] (2-12) |
| **Impact Factor <sup>†</sup>** | 0.84 [0.37-2.29] (0.28-4) |
| **SjR <sup>‡</sup>** | 0.17 [0.13-0.23] (0.1-1.41) |
| **SNIP <sup>§</sup>** | 0.4 [0.23-0.65] (0-4.56) |
| **CiteScore <sup>°</sup>** | 0.32 [0.17-0.46] (0-10.33) |
| **SCImago Quartile** |  |
| **Q1 (% , n)** | 3.3 (10/299) |
| **Q2 (% , n)** | 9.7 (29/299) |
| **Q3 (% , n)** | 40.8 (122/299) |
| **Q4 (% , n)** | 46.1 (138/299) |

Data are reported as medians, interquartile ranges [IQRs] and ranges (minimum value – maximum value) or as percentages and fractions.
\* No missing data. The analyses were conducted on all the 317 Scopus discontinued journals.
<sup>†</sup> Data were available and calculated for 10 journals.
<sup>‡</sup> Data were available and calculated for 304 journals.
<sup>§</sup> Data were available and calculated for 299 journals.
° Data were available and calculated for 82 journals. SJR: SCImago Journal & Country Rank; SNIP: Source Normalized Impact *per* Paper; IF: Impact Factor

### Citation metrics

**Table 2** shows the total number of documents and citations, the total number of documents per journal and the citations count before and after Scopus discontinuation. The total number of citations received after discontinuation was 607,261, with a median of 713 citations (IQR 254-2,056, range 0-19,468) per journal.

**Table 2.** Citations and documents before and after Scopus discontinuation.

|  |  |  |
| --- | --- | --- |
| **Total number of documents** | 591968 |  |
| **Total number of citations** | 1191885 |  |
| **Documents *per* journal\*** | 429 [159.5-1244] (2-132482) |  |
|  | **Before Scopus**<br>**discontinuation** | **After Scopus**<br>**discontinuation** |
| **Citations (n)** | 584624 | 607621 |
| **Citations *per* journal\*** | 415 [120-1580] (0-67529) | 713 [254-2056] (0-19468) |
| **Citations *per* year\*** | 60.32 [17.98-168] (0-4828) | 152.9 [49.43-408] (0-4571) |
| **Citations *per* document\*** | 1 [0.39-2.15] (0-170.4) | 1.66 [0.93-2.66] (0-80.70) |

Data are reported as medians, interquartile ranges [IQRs] and ranges (minimum val- ue – maximum value), if not otherwise specified.
\* No missing data. The analyses were conducted on all the 317 Scopus discontinued journals.

Paired *t*-tests revealed that the mean number of citations per year after discontinuation was significantly higher than before (median of difference 64 citations, p<0.0001). Likewise, the number of citations per document proved significantly higher after discontinuation (median of difference 0.4 citations, p<0.0001) (Table 2).

### Indexing in Cabell’s lists, updated Beall’s list, DOAJ and scientific databases

Twenty-two percent (72/317) of the journals were included in the Cabell’s blacklist, while 29 (9%) were currently under review for inclusion. Only five journals (2%) were in included in the Cabell’s whitelist. In 243 cases (243/317), either the journal’s publisher was included in the updated Beall’s list of predatory publishers or the journal was included in the corresponding list of standalone journals (76.6%). The DOAJ currently includes only 9 journals. Sixty-one journals were previously included and discontinued by DOAJ; in 36 cases the reason was ‘suspected editorial misconduct by the publisher’ while in 23 instances it was ‘journal not adhering to best practice’ and in one case ‘no open access or license info’.

**Table 3** shows the indexing in PubMed and Web of Science.

**Table 3.**
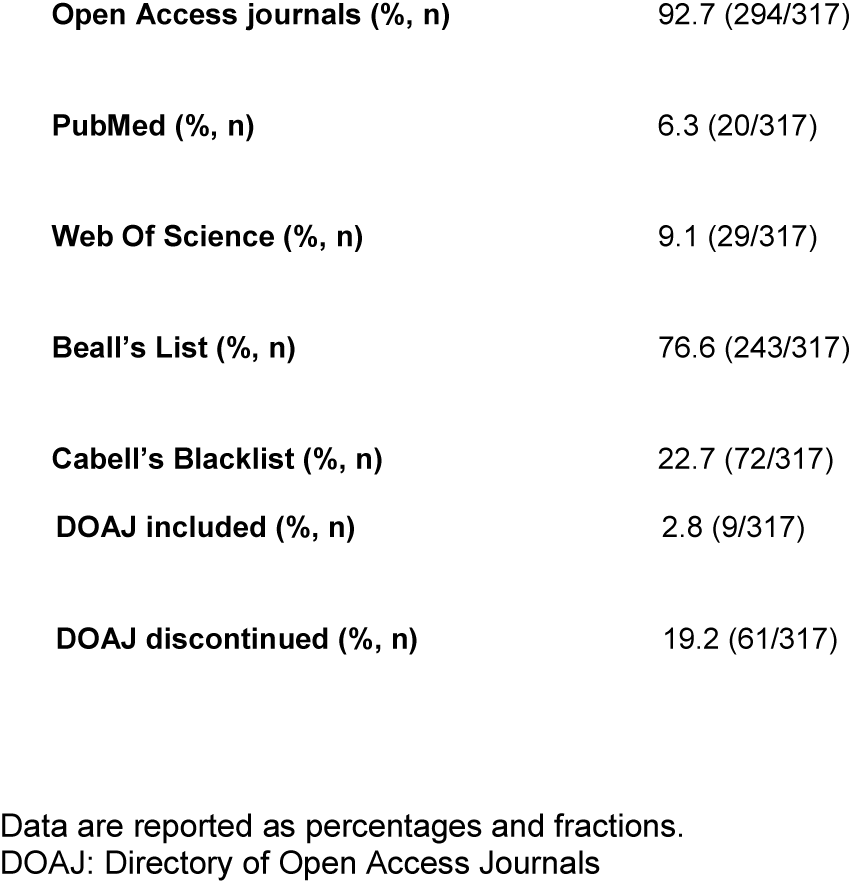
Discontinued journals current Open Access policy and main data-bases indexing.

## Discussion

The present study was aimed at scrutinizing the main features of journals whose coverage was discontinued by Scopus due to publication concerns. To do so, (a) we counted and compared citation metrics per journal and per document obtained *before* and *after* discontinuation, and (b) accessed well-known and established blacklists and whitelists dealing with the issue of predatory publishing, i.e. Cabell’s and updated Beall’s list, as well as the DOAJ. Our main finding was that articles published in these journals before discontinuation, remain available to users and continue to receive a relevant number of citations after discontinuation, more than before. Moreover, a large number of the discontinued journals are likely to be predatory.

Although Scopus applies a rigorous control of content quality and warns users when a journal is discontinued in its source details, the average users tend not to access journal’s details but articles’ contents. By doing so, they remain unaware that the article they have accessed was issued by a journal discontinued for publication concerns. As a consequence, articles issued by journals whose scientific reputation is currently deemed questionable, continue to be displayed and to get cited as contents from legitimate, up-to-standard journals. When quantifying how coverage discontinuation affected the likelihood of these journals to be cited, data indicate that their articles received significantly more citations after discontinuation than before.

Beyond the dangerous exposure of scholars, clinicians and even patients to potentially dubious or low quality contents, the considerable number of citations received after discontinuation by “ghost journals” can be a serious threat to scientific quality assessment by institutions and academia. In fact, these citations contribute to the calculation of the authors’ metrics by Scopus, including the Hirsch index (H-index), [24] which is still among the main descriptors of productivity and scientific impact, based on career advancements are determined. [2,3,4] The fact that “ghost journals” can help to move up in academia is a relevant issue, and has inspired the allegorical vignette depicted in **Fig. 1**: ghost journals can inflate authors’ metrics lifting them unnaturally and effortlessly.

Of greatest concern is our finding that many of the discontinued journals display predatory behaviors in claiming to be open access. Exploitation of the open-access publishing model has been shown to go hand in hand with deviation from best editorial and publication practices for self-interest. [9] Such journals are not only associated with poor editorial quality, but are also deceptive and misleading by nature, i.e. they prioritize self-interest at the expense of scholars, and lack transparent and independent peer review. [9,25] Young researchers from low-income and middle-income countries are probably most susceptible to the false promises and detrimental practices of predatory journals. However, “predatory scholars” also seem to exist, possibly sharing a common interest with deceptive journals and publishers and knowingly using them to achieve their own ends. [7,26,27]

The policy underlying the decision to keep publications prior to discontinuation of indexing is clear. Some of these publications may actually fulfill publishing criteria (e.g. International Committee of Medical Journal Editors, Committee on Publication Ethics). It would be unfair to punish researchers for an eventual deterioration in journal performance; changes in the standards employed by the journal may change over time and the researchers may be unaware of quality issues. On the other hand, as the integrity of the editorial process cannot be vouched for, it is ethically untenable to keep such data available without clearer warnings.

We believe that Scopus should evaluate deleting the discontinued journals from the database contents or, at least, stop tracking their citations. In alternative, we propose that the CSAB could apply these measures case-by-case, after evaluating the severity of the potential misconducts. At the author-level, an alternative may be the provision of two metrics: one with and one without citations from publications in discontinued journals.

This analysis is not free of limitations. First, we included the year of discontinuation in the “*after discontinuation”* period, starting from January 1^th^. This decision may have led to some overestimation in the number of citations received after discontinuation. Second, we included only those journals discontinued from Scopus for “publication concerns” but were not able to retrieve details regarding the specific concern raised. Finally, we did not evaluate the impact of the citations received after discontinuation on author-level metrics.

## Conclusions

The citation count of journals whose coverage in Scopus has been halted for publication concerns, increases despite discontinuation. This paradoxical trend can inflate scholars’ metrics prompting career advancements and promotions. Counter-measures should be taken to ensure the validity and reliability of Scopus metrics both at journals- and author-level for the purpose of scientific assessment of scholarly publishing. Creative thinking is required to resolve this issue without punishing authors who have inadvertently published good quality papers in a failing or predatory journal.

## Acknowledgments

We would like to thank Dr. Antonio Corrado (“Korrado 20”) for creating and providing the **Fig. 1**.

**Figure 1.**
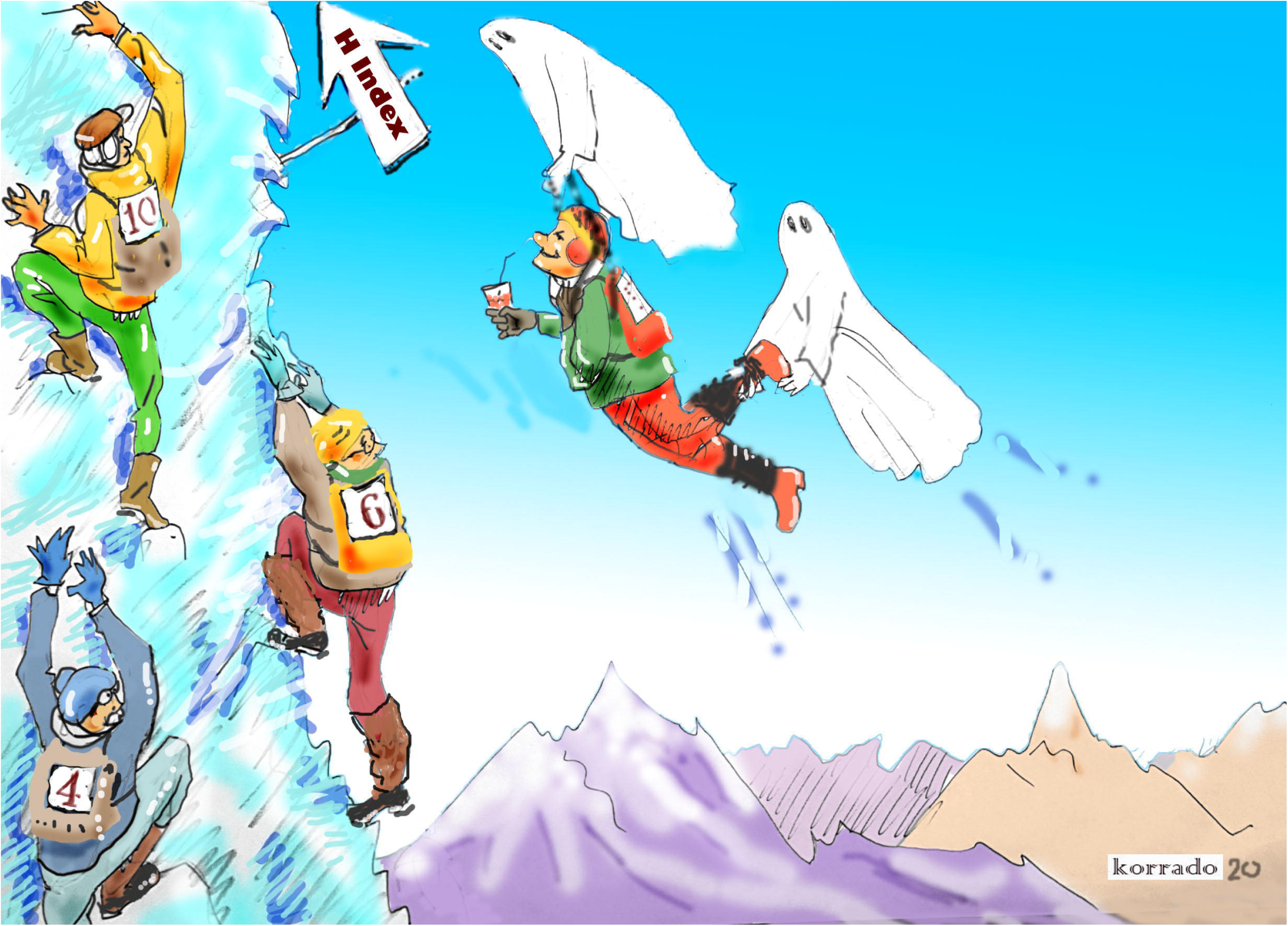
Ghost journals can inflate authors’ metrics lifting them unnaturally and effortlessly.

## Author contributions

**Conceptualization:** Andrea Cortegiani

**Data Curation:** Andrea Cortegiani, Mariachiara Ippolito, Giulia Ingoglia, Andrea Manca, Lucia Cugusi, Anna Severin, Martina Strinzel, Vera Panzarella

**Formal Analysis**: Andrea Cortegiani, Mariachiara Ippolito, Giulia Ingoglia

**Supervision:** Giuseppina Campisi, Lalu Manoj, Cesare Gregoretti, Sharon Einav, David Moher and Antonino Giarratano

**Writing – Original Draft Preparation:** Andrea Cortegiani, Mariachiara Ippolito, Giulia Ingoglia, Andrea Manca, Lucia Cugusi, Anna Severin, Martina Strinzel, **Writing – Review & Editing:** Andrea Cortegiani, Mariachiara Ippolito, Giulia Ingoglia, Andrea Manca, Lucia Cugusi, Anna Severin, Martina Strinzel, Vera Panzarella, Giuseppina Campisi, Lalu Manoj, Cesare Gregoretti, Sharon Einav, David Moher and Antonino Giarratano

