## Supplementary material for "Inflated citations and metrics of journals discontinued from Scopus for publication concerns: the GhoS(t)copus Project": S3 Appendix

**the GhoS(t)copus Project**

Andrea Cortegiani^a*^, Mariachiara Ippolito^a^, Giulia Ingoglia^a^, Andrea Manca^b^, Lucia Cugusi^b^, Anna Severin^c,g^, Michaela Strinzel^c^, Vera Panzarella^d^, Giuseppina Campisi^d^, Lalu Manoj^e^, Cesare Gregoretti^a^, Sharon Einav^f^,

David Moher^e^ and Antonino Giarratano^a^

*^a^Department of Surgical, Oncological and Oral Science (Di.Chir.On.S). Section of Anaesthesia, Analgesia, Intensive Care and Emergency. Policlinico Paolo Giaccone. University of Palermo,* via del Vespro 129, 90127, Palermo, *Italy.*

*^b^Department of Biomedical Sciences, University of Sassari, viale S. Pietro 07100, Sassari, Italy*

*^c^ Swiss National Science Foundation,* *Wildhainweg 3P.O. Box CH-3001 Bern, Switzerland*

*^d^ Department of Surgical, Oncological and Oral Science (Di.Chir.On.S), Section of Oral Medicine, University of Palermo,* via del Vespro 129, 90127, *Palermo, Italy*

*^e^Centre for Journalology, Clinical Epidemiology Program, Ottawa Hospital Research Institute; School of Epidemiology and Public Health, University of Ottawa,* *501 Smyth Road, PO BOX 201B, Ottawa, Ontario, Canada.*

*^f^* *Intensive Care Unit of the Shaare Zedek Medical Medical Centre and Hebrew University Faculty of Medicine, 12 Beyt St, Jerusalem, Israel*

*^g^ Institute for Social and Preventive Medicine, Mittelstrasse 43, 3012, Bern, Switzerland*

*^*^* **Corresponding author**

 (AC)

***Sources***

**Beall's List**

The Beall’s list is a list of “potential, possible, or probable” predatory open access (OA) publishers, created by Jeffrey Beall. It was previously available on Beall’s blog [1], Scholarly Open Access, and now it is kept accessible online by his anonymous successors as an updated online version [2]. Criteria to suspect about a publisher and include it in the list were also provided in details in the blog [3].

**Cabell’s Scholarly Analytics**

Cabell’s Scholarly Analytics is a scholarly, for-profit service that hosts and maintains a whitelist and a blacklist of journals [4]. The whitelist and the blacklist were downloaded in December 2018 and contained 11,057 and 10,671 journals, respectively.

**Whitelist:** the whitelist is intended to provide academics with accurate information and reputable outlets for publication [5]. To be indexed, journals must fulfil a number of criteria, which cover aspects such as Business Practices, Policies, Peer Review or Editorial Services [6].

**Blacklist:** the aim of the blacklist is to warn authors against journals with questionable practices and entails journals that have exhibited behaviour that can be indicative of deception [4]. Journals are checked against more than 60 criteria, which relate to aspects such as Business Practices, Publishing, Archiving and Access and Editorial Services [6, 7].

**Centre for Science and Technology Studies (CWTS)**

CWTS Journal Indicators offers free access to bibliometric indicators on scientific journals, calculated by Leiden University’s CWTS, based on the Scopus database [8].

**Directory of Open Access Journals (DOAJ)**

The DOAJ is a community-curated list of OA journals. The DOAJ is a community of OA publishers with more than 100 volunteers and a core team of 15 people, employed by DOAJ's holding company, IS4OA [6]. DOAJ was launched in 2003 at Lund University in Sweden with 300 OA journals indexed in the very beginning, the number of journals included in DOAJ has already increased dramatically and it contains more than 14,000 OA journals covering all areas of science, technology, medicine, social sciences and humanities [9].

**DOAJ removed titles**

DOAJ contents are regularly reviewed and under certain circumstances, journals can be removed from the DOAJ. Potential reasons for removal can be that a journal is no longer OA, if there is evidence for editorial misconduct or if a journal does not adhere to Best practice [10]. To foster transparency, the DOAJ publishes the list of journals that were removed from their directory. The list of journals removed starts from the beginning of 2014 and includes information on journal title, ISSN, date removed and the reason [10]. As of February 2020, this list included 3482 journals.

**Journal Citation Reports**

Journal Citation Reports collects metrics and analysis of impactful journals included in the *Science Citation Index Expanded (SCIE)* and *Social Sciences Citation Index (SSCI)*, part of the *Web of Science Core Collection*. Journals with evidence of excessive self-citation and citation stacking are usually excluded from Journal Citations Reports [11].

**PubMed**

PubMed is developed and maintained by the *National Center for Biotechnology Information* (NCBI). It contains citations and abstracts mainly from biomedicine and health journals, but it also covers other fields [12].

**SCImago Journal & Country Rank**

SCImago Journal & Country Rank is a public portal containing journals and country scientific indicators developed from the information contained in the Scopus database. SCImago is a research group from the Consejo Superior de Investigaciones Científicas (CSIC), University of Granada, Extremadura, Carlos III (Madrid) and Alcalá de Henares [13].

**Scopus database**

The Scopus database is an abstracts and citations database of peer-reviewed research literature with contents from over 5000 Publishers [14]. Its content is periodically reviewed and selected by an independent *Content Selection and Advisory Board* (CSAB) [14]. It provides also analytic tools, with bibliometric and historic trend analysis.

**Web of Science**

It is a publisher-independent global citation database [15]. Content is selected by a team of editors, in charge of evaluating its quality and impact.

***Metrics***

**CiteScore**

It is a Scopus metric, calculated as the ratio between the number of citations recorded from all documents in year one and the number of documents published in the prior three years [16].

**Impact factor (IF)**

In any given year, the IF of a journal is the number of citations, received in that year, of articles published in that journal during the two preceding years, divided by the total number of "citable items" published in that journal during the two preceding years [17].

**SCImago Journal Rank (SJR) indicator**

SCImago Journal Rank (SJR) indicator, developed by SCImago from the algorithm Google PageRank™, is a measure of scientific influence of scholarly journals that accounts for both the number of citations received by a journal and the importance or prestige of the journals where such citations come from [17]. A journal's SJR indicates the average number of weighted citations received during a selected year per document published in that journal during the previous three years [17].

**Source Normalized Impact per Paper (SNIP)**

Source Normalized Impact per Paper intrinsically accounts for field-specific differences in citation practices. It does so by comparing each journal's citations per publication with the citation potential of its field, defined as the set of publications citing that journal [18].
