## Supplementary material for "Inflated citations and metrics of journals discontinued from Scopus for publication concerns: the GhoS(t)copus Project": S1 Fig, S1 Table, S2 Table, S3 Table, S4 Table

**the GhoS(t)copus Project**

Andrea Cortegiani^a*^, Mariachiara Ippolito^a^, Giulia Ingoglia^a^, Andrea Manca^b^, Lucia Cugusi^b^, Anna Severin^c,g^, Michaela Strinzel^c^, Vera Panzarella^d^, Giuseppina Campisi^d^, Lalu Manoj^e^, Cesare Gregoretti^a^, Sharon Einav^f^,

David Moher^e^ and Antonino Giarratano^a^

*^a^Department of Surgical, Oncological and Oral Science (Di.Chir.On.S). Section of Anaesthesia, Analgesia, Intensive Care and Emergency. Policlinico Paolo Giaccone. University of Palermo,* via del Vespro 129, 90127, Palermo, *Italy.*

*^b^Department of Biomedical Sciences, University of Sassari, viale S. Pietro 07100, Sassari, Italy*

*^c^ Swiss National Science Foundation,* *Wildhainweg 3P.O. Box CH-3001 Bern, Switzerland*

*^d^ Department of Surgical, Oncological and Oral Science (Di.Chir.On.S), Section of Oral Medicine, University of Palermo,* via del Vespro 129, 90127, *Palermo, Italy*

*^e^Centre for Journalology, Clinical Epidemiology Program, Ottawa Hospital Research Institute; School of Epidemiology and Public Health, University of Ottawa,* *501 Smyth Road, PO BOX 201B, Ottawa, Ontario, Canada.*

*^f^* *Intensive Care Unit of the Shaare Zedek Medical Medical Centre and Hebrew University Faculty of Medicine, 12 Beyt St, Jerusalem, Israel*

*^g^ Institute for Social and Preventive Medicine, Mittelstrasse 43, 3012, Bern, Switzerland*

**CONTENT**

**Page 3, S1 Table. Distribution of Scopus discontinued journals by Publisher**

**Page 4, S1 Fig. and S2 Table. Distribution of Scopus discontinued journals by country**

**Page 5, S3 Table. Distribution of Scopus discontinued journals by *subject areas***

**Page 6, S4 Table. Distribution of Scopus discontinued journals by subject sub-areas.**

| ***Publishers (n=135)*** | ***% ( n )*** |
| --- | --- |
| *Academic Journals Inc.*  *Asian Network for Scientific Information*  *OMICS Publishing Group*  *Medwell Journals*  *iMedPub*  *World Scientific and Engineering Academy and Society*  *Science Publications*  *Academy Publisher* | 12.3 (39/317)  6 (19/317)  5.7 (18/317)  4.1 (13/317)  3.5 (11/317)  2.8 (9/317)  2.5 (8/317)  2.2 (7/317) |
| *Allied Academies*  *Canadian Center of Science and Education*  *International Digital Organization for Scientific Information (IDOSI)* | 1.9 (6/317)  1.9 (6/317)  1.9 (6/317) |
| *Science and Engineering Research Support Society*  *Serials Publications (International Science Press)*  *AMSE Press*  *Eurojournals Inc.*  *Hikari Ltd*  *Research India Publications*  *Others* | 1.6 (5/317)  1.6 (5/317)  1.3 (4/317)  1.3 (4/317)  1.3 (4/317)  1.3 (4/317)  47 (149/317) |

**S1 Table. Distribution of Scopus discontinued journals by publisher.**

Data are reported as percentages and fractions. Publishers with less than four Scopus discontinued journals were grouped as ‘Others’.

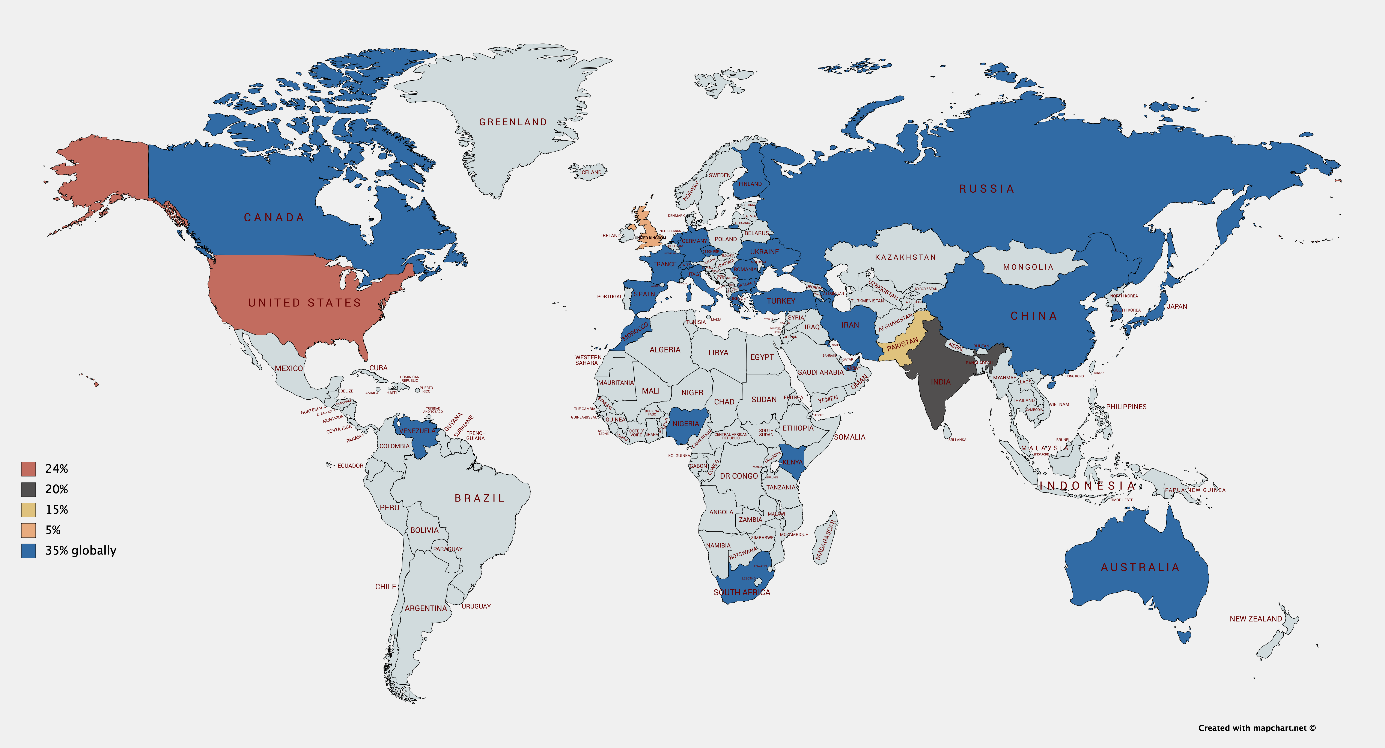

**S1 Fig. Distribution of Scopus discontinued journals by country.**

The map chart shows the different frequencies of distribution by country with different colors.

| ***Country (n=33)*** | ***% ( n )*** |
| --- | --- |
| *United States of America*  *India*  *Pakistan* | 24 (76/316)  19.9 (63/316)  15.5 (49/316) |
| *United Kingdom* | 5.4 (17/316) |
| *Turkey* | 4.1 (13/316) |
| *Greece* | 3.5 (11/316) |
| *Canada* | 3.2 (10/316) |
| *Finland* | 2.5 (8/316) |
| *France*  *United Arab Emirates*  *Italy*  *Romania*  *South Korea*  *Ukraine*  *Australia*  *Bulgaria*  *Others* | 2.2 (7/316)  1.9 (6/316)  1.6 (5/316)  1.6 (5/316)  1.6 (5/316)  1.6 (5/316)  1.3 (4/316)  1.3 (4/316)  8.8 (28/316) |

**S2 Table. Distribution of Scopus discontinued journals by country.**

Data were retrieved from Scimago Journal & Country Rank and are reported as percentages and fractions. Countries with less than four Scopus discontinued journals were grouped as ‘Others’

| Medicine | 16.40% | (52/317) |
| --- | --- | --- |
| Agricultural and Biological Sciences | 10.70% | (34/317) |
| Pharmacology, Toxicology and Pharmaceutics | 9.77% | (31/317) |
| Engineering | 7.88% | (25/317) |
| Computer Science | 7.88% | (25/317) |
| Biochemistry, Genetics and Molecular Biology | 5.36% | (18/317) |
| Business, Management and Accounting | 5.36% | (17/317) |
| Mathematics | 5.36% | (17/317) |
| Social Sciences | 4.73% | (15/317) |
| Arts and Humanities | 3.78% | (12/317) |
| Multidisciplinary | 3.50% | (11/317) |
| Economics, Econometrics and Finance | 2.52% | (8/317) |
| Environmental Science | 2.21% | (7/317) |
| Immunology and Microbiology | 2.21% | (7/317) |
| Materials Science | 2.21% | (7/317) |
| Veterinary | 2.21% | (7/317) |
| Earth and Planetary Sciences | 1.58% | (5/317) |
| Chemistry | 1.26% | (4/317) |
| Energy | 1.26% | (4/317) |
| Chemical Engineering | 0.94% | (3/317) |
| Physics and Astronomy | 0.94% | (3/317) |
| Nursing | 0.63% | (2/317) |
| Dentistry | 0.32% | (1/317) |
| Health Professions | 0.32% | (1/317) |
| Neuroscience | 0.32% | (1/317) |

**S3 Table. Distribution of Scopus discontinued journals by subject areas.**

Data were retrieved from Scopus and are reported as percentages and fractions.

| **Medicine** | 16.40% | (52/317) | General Medicine | 21.15% | (11/52) |
| --- | --- | --- | --- | --- | --- |
|  |  |  | Complementary and Alternative Medicine | 11.54% | (6/52) |
|  |  |  | Oncology | 11.54% | (6/52) |
|  |  |  | Infectious Diseases | 9.61% | (5/52) |
|  |  |  | Cardiology and Cardiovascular Medicine | 5.77% | (3/52) |
|  |  |  | Pharmacology (medical) | 5.77% | (3/52) |
|  |  |  | Pediatrics, Perinatology and Child Health | 3.84% | (2/52) |
|  |  |  | Immunology and Allergy | 3.84% | (2/52) |
|  |  |  | Psychiatry and Mental Health | 3.84% | (2/52) |
|  |  |  | Genetics (clinical) | 1.92% | (1/52) |
|  |  |  | Anesthesiology and Pain Medicine | 1.92% | (1/52) |
|  |  |  | Biochemistry (medical) | 1.92% | (1/52) |
|  |  |  | Epidemiology | 1.92% | (1/52) |
|  |  |  | Geriatrics and Gerontology | 1.92% | (1/52) |
|  |  |  | Health Policy | 1.92% | (1/52) |
|  |  |  | Internal Medicine | 1.92% | (1/52) |
|  |  |  | Medicine (Miscellaneus) | 1.92% | (1/52) |
|  |  |  | Obstetrics and Gynecology | 1.92% | (1/52) |
|  |  |  | Otorhinolaryngology | 1.92% | (1/52) |
|  |  |  | Public Health, Enviromental and Occupational Health | 1.92% | (1/52) |
|  |  |  | Pulmonary and Respiratory Medicine | 1.92% | (1/52) |
| **Agricultural and Biological Sciences** | 10.70% | (34/317) | Agronomy and Crop Science | 23.52% | (8/34) |
|  |  |  | Food Science | 17.65% | (6/34) |
|  |  |  | General Agricultural and Biological Sciences | 14.70% | (5/34) |
|  |  |  | Animal Science and Zoology | 11.76% | (4/34) |
|  |  |  | Horticulture | 8.82% | (3/34) |
|  |  |  | Plant Science | 8.82% | (3/34) |
|  |  |  | Agricultural and Biological Sciences (miscellaneous) | 2.50% | (1/34) |
|  |  |  | Aquatic Science | 2.50% | (1/34) |
|  |  |  | Ecology, Evolution, Behavior and Systematics | 2.50% | (1/34) |
|  |  |  | Forestry | 2.50% | (1/34) |
|  |  |  | Insect Science | 2.50% | (1/34) |
| **Pharmacology, Toxicology and Pharmaceutics** | 9.77% | (31/317) | Pharmaceutical Science | 48.38% | (15/31) |
|  |  |  | General Pharmacology, Toxicology and Pharmaceutics | 32.25% | (10/31) |
|  |  |  | Drug Discovery | 9.67% | (3/31) |
|  |  |  | Pharmacology | 9.67% | (1/31) |
| **Engineering** | 7.88% | (25/317) | General Engineering | 40% | (10/25) |
|  |  |  | Electrical and Electronic Engineering | 24% | (6/25) |
|  |  |  | Control and Systems Engineering | 16% | (4/25) |
|  |  |  | Ocean Engineering | 12% | (3/25) |
|  |  |  | Industrial and Manufacturing Engineering | 4% | (1/25) |
|  |  |  | Mechanical Engineering | 4% | (1/25) |
| **Computer Science** | 7.88% | (25/317) | General Computer Science | 44% | (11/25) |
|  |  |  | Computer Networks and Communications | 16% | (4/25) |
|  |  |  | Artifical Intelligence | 12% | (3/25) |
|  |  |  | Computer Science Applications | 12% | (3/25) |
|  |  |  | Software | 12% | (3/25) |
|  |  |  | Information Systems | 4% | (1/25) |
| **Biochemistry, Genetics and Molecular Biology** | 5.36% | (18/317) | Biochemistry | 22.20% | (4/18) |
|  |  |  | General Biochemistry, Genetics and Molecular Biology | 22.20% | (4/18) |
|  |  |  | Biotechnology | 16.67% | (3/18) |
|  |  |  | Cancer Research | 11.11% | (2/18) |
|  |  |  | Biochemistry, Genetics and Molecular Biology (miscellaneous) | 5.55% | (1/18) |
|  |  |  | Cell Biology | 5.55% | (1/18) |
|  |  |  | Developmental Biology | 5.55% | (1/18) |
|  |  |  | Endocrinology | 5.55% | (1/18) |
|  |  |  | Molecular Medicine | 5.55% | (1/18) |
| **Business, Management and Accounting** | 5.36% | (17/317) | Business and International Management | 41.18% | (7/17) |
|  |  |  | General Business, Management and Accounting | 35.28% | (6/17) |
|  |  |  | Marketing | 11.76% | (2/17) |
|  |  |  | Management Information Systems | 5.90% | (1/17) |
|  |  |  | Strategy and Management | 5.90% | (1/17) |
| **Mathematics** | 5.36% | (17/317) | General Mathematics | 41.18% | (7/17) |
|  |  |  | Applied Mathematics | 17.65% | (3/17) |
|  |  |  | Modeling and Simulation | 17.65% | (3/17) |
|  |  |  | Algebra and Number Theory | 11.76% | (2/17) |
|  |  |  | Analysis | 5.88% | (1/17) |
|  |  |  | Computational Mathematics | 5.88% | (1/17) |
| **Social Sciences** | 4.73% | (15/317) | Education | 26.66% | (4/15) |
|  |  |  | General Social Sciences | 20% | (3/15) |
|  |  |  | Anthropology | 13.33% | (2/15) |
|  |  |  | Communication | 13.33% | (2/15) |
|  |  |  | Gender Studies | 6.66% | (1/15) |
|  |  |  | Health (social science) | 6.66% | (1/15) |
|  |  |  | Safety Research | 6.66% | (1/15) |
|  |  |  | Sociology and Political Sciences | 6.66% | (1/15) |
| **Arts and Humanities** | 3.78% | (12/317) | Language and Linguistics | 33.30% | (4/12) |
|  |  |  | General Arts and Humanities | 25% | (3/12) |
|  |  |  | Literature and Literary Theory | 25% | (3/12) |
|  |  |  | History | 16.70% | (2/12) |
| **Multidisciplinary** | 3.50% | (11/317) |  |  |  |
| **Economics, Econometrics and Finance** | 2.52% | (8/317) | General Economics, Econometrics and Finance | 62.50% | (5/8) |
|  |  |  | Economics and Econometrics | 37.50% | (3/8) |
| **Environmental Science** | 2.21% | (7/317) | General Environmental Science | 42.85% | (3/7) |
|  |  |  | Ecology | 14.28% | (1/7) |
|  |  |  | Pollution | 14.28% | (1/7) |
|  |  |  | Waste Management and Disposal | 14.28% | (1/7) |
|  |  |  | Water Science and Technology | 14.28% | (1/7) |
| **Immunology and Microbiology** | 2.21% | (7/317) | Immunology | 42.85% | (3/7) |
|  |  |  | Applied Microbiology and Biotechnology | 28.57% | (2/7) |
|  |  |  | Virology | 28.57% | (2/7) |
| **Materials Science** | 2.21% | (7/317) | General Materials Science | 42.85% | (3/7) |
|  |  |  | Ceramics and Composites | 14.28% | (1/7) |
|  |  |  | Materials Chemistry | 14.28% | (1/7) |
|  |  |  | Metals and Alloys | 14.28% | (1/7) |
|  |  |  | Polymers and Plastics | 14.28% | (1/7) |
| **Veterinary** | 2.21% | (7/317) | General Veterinary | 85.71% | (6/7) |
|  |  |  | Food Animals | 14.28% | (1/7) |
| **Earth and Planetary Sciences** | 1.58% | (5/317) | General Earth and Planetary Sciences | 40% | (2/5) |
|  |  |  | Atmospheric Science | 20% | (1/5) |
|  |  |  | Geotechnical Engineering and Engineering Geology | 20% | (1/5) |
|  |  |  | Space and Planetary Science | 20% | (1/5) |
| **Chemistry** | 1.26% | (4/317) | General Chemistry | 50% | (2/4) |
|  |  |  | Physical and Theoretical Chemistry | 50% | (2/4) |
| **Energy** | 1.26% | (4/317) | General Energy | 75% | (3/4) |
|  |  |  | Renewable Energy, Sustainability and the Environment | 25% | (1/4) |
| **Chemical Engineering** | 0.94% | (3/317) | General Chemical Engineering | 66.66% | (2/3) |
|  |  |  | Bioengineering | 33.33% | (1/3) |
| **Physics and Astronomy** | 0.94% | (3/317) | General Physics and Astronomy | 100% | (3/3) |
| **Nursing** | 0.63% | (2/317) | General Nursing | 50% | (1/2) |
|  |  |  | Nutrition and Dietetics | 50% | (1/2) |
| **Dentistry** | 0.32% | (1/317) | General Dentistry | 100% | (1/1) |
| **Health Professions** | 0.32% | (1/317) | Physical Therapy, Sports Therapy and Rehabilitation | 100% | (1/1) |
| **Neuroscience** | 0.32% | (1/317) | General Neuroscience | 100% | (1/1) |

**S4 Table. Distribution of Scopus discontinued journals by subject sub-areas.**

Data were retrieved from Scopus and are reported as percentages and fractions.
